# An Integrated Risk Predictor for Pulmonary Nodules

**DOI:** 10.1101/094920

**Authors:** Paul Kearney, Xiao-Jun Li, Alex Porter, Steve Springmeyer, Peter Mazzone

**Affiliations:** Integrated Diagnostics, Seattle, WA; Respiratory Institute, The Cleveland Clinic, Cleveland, OH

**Keywords:** Pulmonary nodules, Proteomic, Biomarker, Lung cancer, Risk prediction

## Abstract

It is estimated that over 1.5 million lung nodules are detected annually in the United States. Most of these are benign but frequently undergo invasive and costly procedures to rule out malignancy. A risk predictor that can accurately differentiate benign and malignant lung nodules could be used to more efficiently route benign lung nodules to non-invasive observation by CT surveillance and route malignant lung nodules to invasive procedures. The majority of risk predictors developed to date are based exclusively on clinical risk factors, imaging technology or molecular markers. Assessed here are the relative performances of previously reported clinical risk factors and proteomic molecular markers for assessing cancer risk in lung nodules. From this analysis an integrated model incorporating clinical risk factors and proteomic molecular markers is developed and its performance assessed on a previously reported prospective collection of lung nodules that enrolled 475 patients from 12 sites with lung nodules between 8 and 30mm in diameter. In this analysis it is found that the molecular marker is most predictive. However, the integration of clinical and molecular markers is superior to both clinical and molecular markers separately.

Clinical Trial Registration: Registered at ClinicalTrials.gov (NCT01752101).

## INTRODUCTION

Over 1.5 million lung nodules are identified annually in the U.S. presenting a difficult clinical challenge as the majority ultimately prove to be of benign origin (*1, 2*). The diagnostic dilemma faced by physicians is to identify nodules that are malignant and yet minimize the risks of invasive procedures on benign nodules. Evidence suggests that the currently available tools such as clinical risk predictors (*3-5*) and imaging have limitations in clinical practice (*6, 7*), This has resulted in a growing interest in utilizing molecular tests as diagnostic adjuncts (*8, 9*). Perhaps of even greater interest is the utility of integrated risk predictors that incorporate both clinical risk factors and molecular markers.

We have previously validated a blood-based risk predictor that used 11 molecular factors (*10,11*) and established its potential clinical utility on a prospectively collected biobank (*12*). Separately, several risk predictors composed purely of clinical risk factors have been validated (*3-5*). Here we explore two hypotheses:

- First, molecular markers are comparable in performance to clinical factors for risk prediction, and
- Second, the integration of molecular and clinical risk factors results in a better risk prediction.

The most reliable clinical tools are those that undergo multiple validations of performance on independent sample sets (*13*). Consequently, the focus of this analysis is not the discovery of new molecular or clinical risk factors, but the evaluation of previously validated markers and factors, both individually and integrated, on a prospectively collected sample set. The molecular markers evaluated are those previously discovered and validated as being predictive for cancer risk in lung nodules (*8,11*). The clinical risk factors, also previously validated as being predictive for cancer risk in lung nodules, include age, nodule size, smoking history, nodule location and nodule spiculation (*5*). Finally, the hypotheses are explored using samples acquired from a prospective trial (NCT01752101) of lung nodule management previously reported (*12*). All subjects in this trial underwent an invasive procedure (biopsy and/or surgery). We chose to focus on these subjects as it allows for the assessment of how accurately risk prediction tools could identify benign lung nodules that undergo unnecessary invasive procedures.

## METHODS

### Study Design

Trial NCT01752101 was a prospective, multicenter, observational trial with retrospective evaluation of the performance of molecular and clinical markers. Patient care was not directed or influenced by the protocol. A blinding protocol was strictly followed. All sites had local or central Institutional Review Board approval.

### Patient Selection

Patients with an indeterminate pulmonary nodule were enrolled at 12 geographically diverse sites in the U.S. Eligible patients were those with a lung nodule between 8-30mm in diameter, minimum 40 years of age, and had recently completed a CT guided needle aspiration (TTNA) or bronchoscopic biopsy with an established diagnosis or scheduled for a surgical lung biopsy. Exclusion criteria included a prior malignancy within 5 years of lung nodule identification or a clinical tumor stage > or = T2, nodal stage > or = N2, or evidence of metastatic disease. All TTNA and bronchoscopy procedures were categorized as either diagnostic (provided a specific malignant or benign pathological diagnosis), or non-diagnostic (the specific etiology of the lung nodule remained unknown). All surgical procedures were categorized into either diagnostic (i.e. no specific prior diagnosis) or therapeutic (i.e. surgery preceded by a TTNA or bronchoscopy that yielded a malignant diagnosis).

Finally, the analysis here focused on those subjects with lung nodules between 8 and 20mm in diameter. These were of particular interest as lung nodules below 20mm are more challenging to classify as malignant or benign

### Data Collection

Blood samples were obtained and processed for proper storage and shipment per protocol (*12*). Data on procedures including bronchoscopy, transthoracic needle biopsy, and surgery were recorded.

### Molecular and Clinical Markers

The assay (Xpresys Lung^™^, Integrated Diagnostics, Seattle, WA) is based on mass spectroscopy MRM-MS as previously described (*8,11,12,14*). It was developed following National Academy of Medicine guidelines (*13*) and monitors 11 proteins. Clinical factors collected included age, smoking status, nodule diameter, nodule spiculation and nodule location.

### Data Analysis

All statistical analyses were performed using Matlab (Mathworks inc., version 8.3.0.532) and MedCalc, version 16.4 (MedCalc Software bvba). Chi-squared analysis of variance testing was used to compare groups, and a p-value of ≤ 0.05 is considered significant. The McMenar test was used to compare performance of different predictors on the same sample set (*15*). Physicians, patients, along with laboratory and statistical personnel were blinded to the results of the protein classifier and clinical information. Continuous and categorical variables were assessed using Mann–Whitney and Fisher’s exact tests, respectively. All confidence intervals are reported as two-sided binomial 95% confidence intervals (CIs).

## RESULTS

A total of 475 subjects were enrolled prospectively from April 2012 to December 2014 in the registered study, NCT01752101. Of these, 50 subjects violated the inclusion/exclusion criteria; 43 additional subjects had a lung cancer other than NSCLC; 26 additional subjects had missing data; and 3 additional subjects violated the blood sample collection protocol. This left 353 patients eligible for analysis (see (*12*) for more details). Of the 353 eligible subjects, 222 had nodule size 8-20mm. Baseline demographics are shown in Table 1. In this population of subjects, the cancer prevalence was 81% (180 out of 222 subjects). The average age of subjects with a malignant nodule (67.1 years) was not significantly different from subjects with a benign nodule (64.8 years). Smoking history binned into the categories of ‘never’, ‘former’ and ‘current’ were similar for subjects with malignant and benign nodules. Nodule size was not significantly different between subjects with a malignant nodule (14.7mm) and subjects with a benign nodule (14.1mm).

**TABLE 1.**
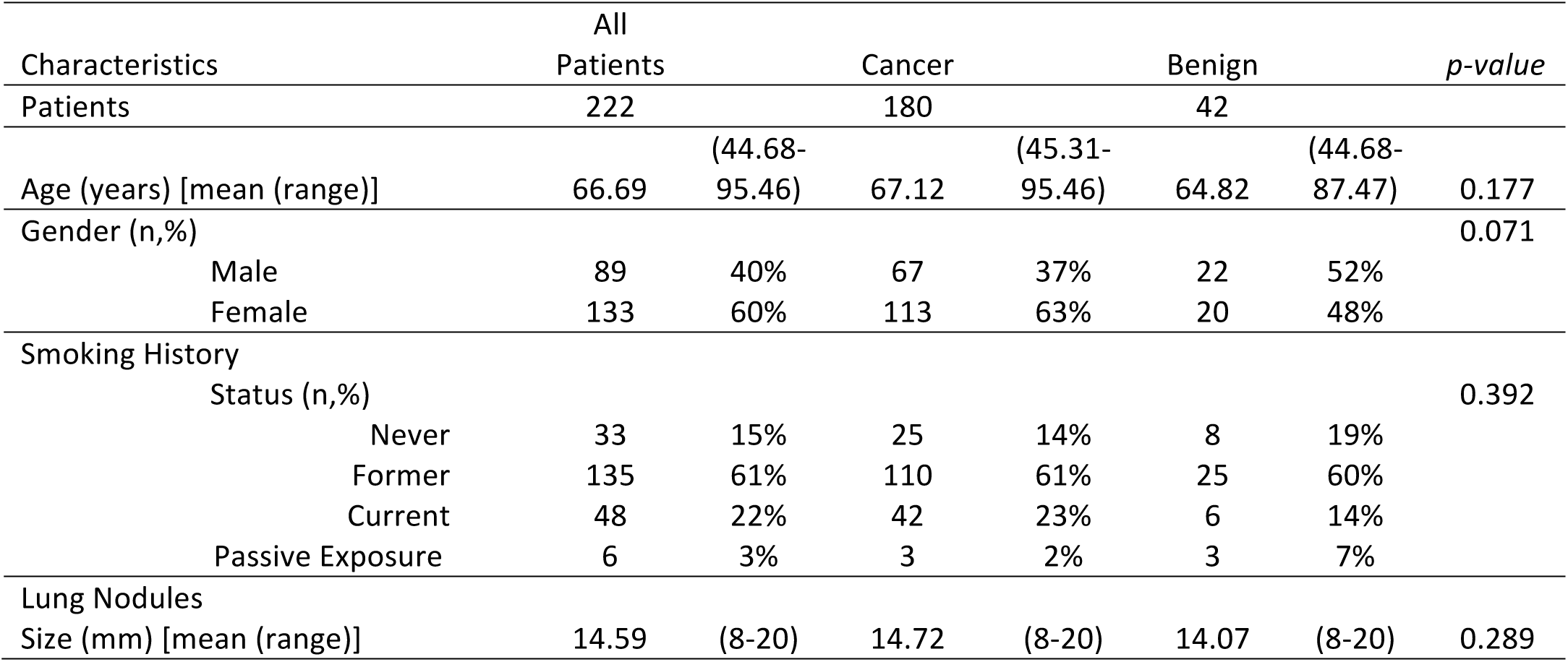
Patient demographics and lung nodule characteristics for all 222 subjects.

Of the 11 proteomic markers evaluated five were previously reported as being diagnostic (ALDOA, COIA1, TSP1, FRIL and LG3BP) and 6 reported as being endogenous normalizers (C163A, PEDF, LUM, GELS, MASP and PTPRJ) (*11,14*), In this analysis, we focus on ratio pairs P1/P2 were P1 is a diagnostic protein and P2 is a normalizer. Among all such protein ratio pairs, LG3BP/C163A had the maximal AUC of 60% for classifying the 222 subjects as being malignant or benign. This is significantly better than random by the Mann-Whitney test (p-value 0.025). The ratio of these two proteins, LG3BP/C163A, will be used to assess the stated study hypotheses. Table 2 presents the AUC performance of all ratio pairs.

**TABLE 2.** AUC performance of all proteomic ratio pairs P1/P2 where P1 is one of the five diagnostic proteins (ALDOA, COIA1, TSP1, FRIL and LG3BP) and P2 is one of the six normalization proteins (C163A, PEDF, LUM, GELS, MASP and PTPRJ).

| P1 | P2 | AUC |
| --- | --- | --- |
| LG3BP | C163A | 0.60 |
| LG3BP | GELS | 0.60 |
| COIA1 | GELS | 0.59 |
| COIA1 | LUM | 0.56 |
| COIA1 | PEDF | 0.56 |
| ALDOA | C163A | 0.56 |
| FRIL | LUM | 0.56 |
| FRIL | PEDF | 0.55 |
| LG3BP | MASP1 | 0.55 |
| LG3BP | PTPRJ | 0.55 |
| ALDOA | GELS | 0.54 |
| FRIL | MASP1 | 0.54 |
| TSP1 | C163A | 0.54 |
| TSP1 | GELS | 0.53 |
| COIA1 | C163A | 0.53 |
| ALDOA | PTPRJ | 0.53 |
| ALDOA | PEDF | 0.53 |
| FRIL | PTPRJ | 0.52 |
| COIA1 | MASP1 | 0.52 |
| TSP1 | LUM | 0.52 |
| TSP1 | PTPRJ | 0.52 |
| TSP1 | PEDF | 0.51 |
| FRIL | GELS | 0.51 |
| LG3BP | PEDF | 0.51 |
| ALDOA | LUM | 0.51 |
| FRIL | C163A | 0.51 |
| COIA1 | PTPRJ | 0.51 |
| TSP1 | MASP1 | 0.50 |
| LG3BP | LUM | 0.50 |
| ALDOA | MASP1 | 0.50 |

### Study Hypothesis #1

We assess the individual performance of five clinical risk factors (nodule size, subject age, subject smoking history, nodule location and nodule spiculation) and the proteomic ratio LG3BP/C163A. Figure 1 presents the performance of the five clinical risk factors and the molecular marker LG3BP/C163A using a ROC plot. In comparing these six factors, the proteomic ratio has the highest AUC (60%) and is significantly better than random (pvalue = 0.025). In contrast, none of the clinical risk factors alone have an AUC significantly better than random (AUC appears in parentheses): nodule size (55%), age (57%), smoking history (55%), nodule location (48%) and nodule spiculation (55%).

**Figure 1.**
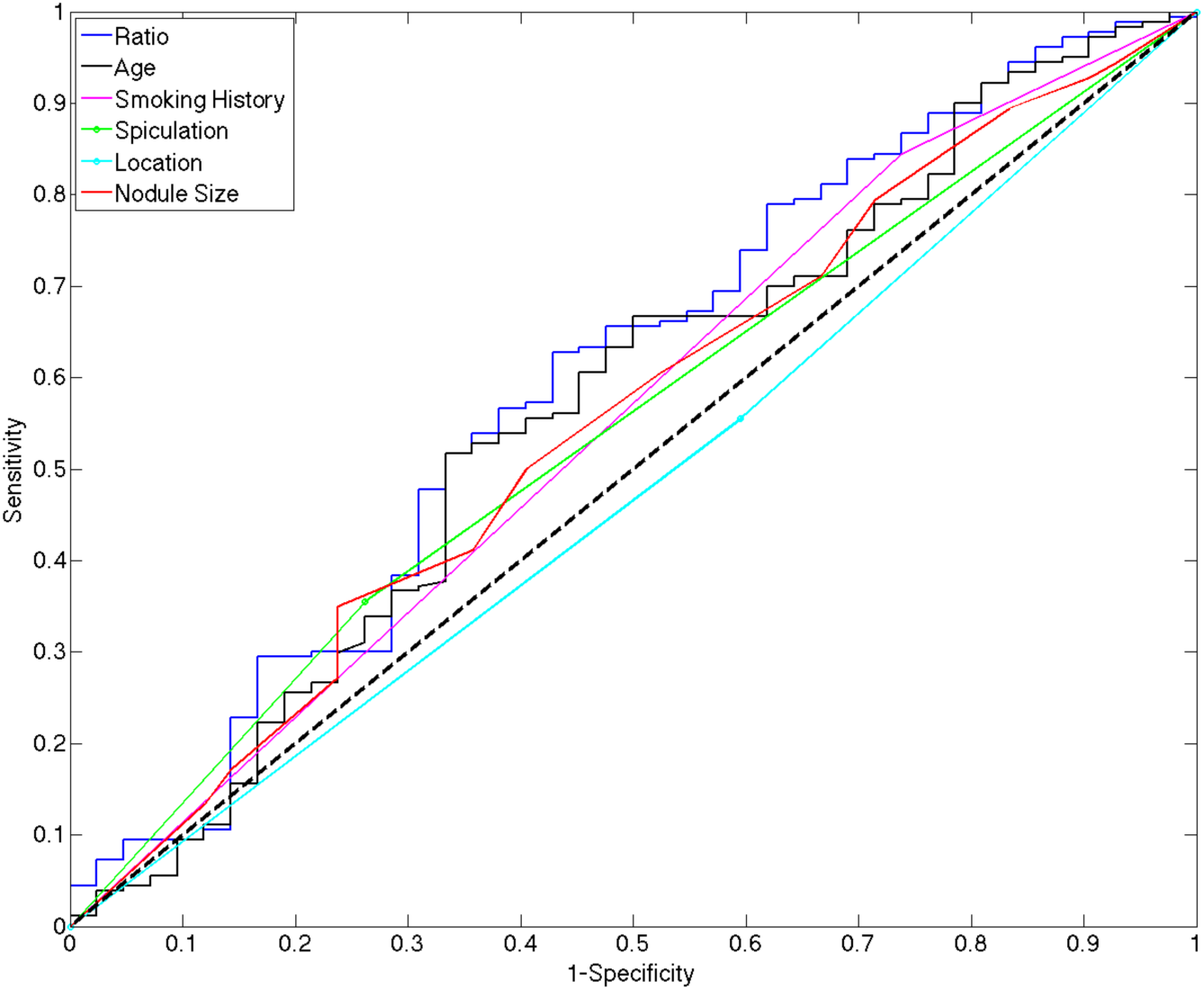
Comparison of five clinical risk factors and the proteomic ratio LG3BP/C163A.

### Study Hypothesis #2

To address the second study hypothesis, we need to integrate the five clinical risk factors with the proteomic ratio. This integration was conducted using a decision tree approach (*16*). The proteomic ratio LG3BP/C163A is used first to classify a lung nodule as lower or higher risk based on a decision threshold “t” (see formula below where “k” denotes a lung nodule under evaluation). Secondly, a cancer risk score, “ClinFact”, is calculated over the five clinical risk factors. Note that the ClinFact risk algorithm is a simplification of the ‘Mayo’ algorithm (*5*) with the cancer history factor omitted as insufficient information was collected in the study for this clinical factor to be reliably included. If the lung nodule is lower risk, as determined by the proteomic ratio LG3BP/C163A, then the Mayo risk prediction is reduced by a fixed amount “T”, otherwise, it is left unchanged. We call this integrated model “IntMod”. Effectively, the proteomic ratio LG3BP/C163A is used to identify lower risk lung nodules in order to rule out lung cancer.

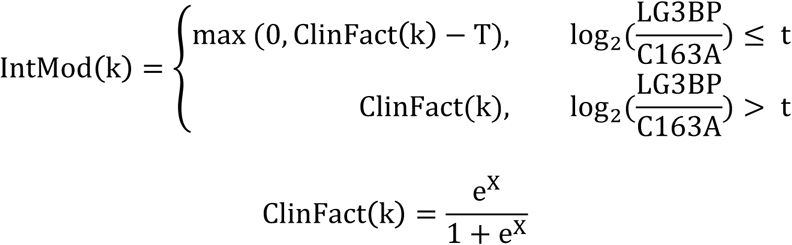

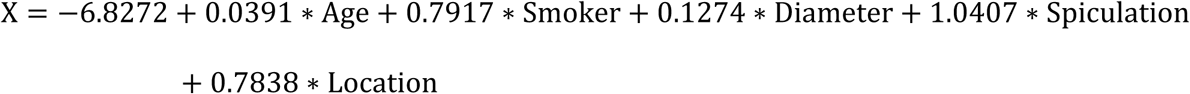

We note that this integrated model has two parameters, namely, “t” and “T” which need to be learned.

### Optimal Values of Parameters t and T

In this analysis we learn the optimal values for the integrating parameters t and T. All possible pairs of values for t (−1.1 to 3.3 in increments of 0.1) and for T (0 to 1 in increments of .1) are assessed and the AUC of the resulting IntMod predictor calculated. We make the following observations:

1. The longest continuous stretch of values for t where the highest AUC values occur is from t = 0. 14 to 0.39, regardless of the value of parameter T. In this range the AUC values are all at least 62%. That is, this is the range of values for parameter t with both high and stable performance for IntMod.
2. Within the range t = 0.14 to 0.39, the vast majority of t values (23 out of 29) achieve maximal AUC when T = 0.5. In particular, the maximum AUC achieved is 63.1% for t = 0.29 and T = 0.5.
3. Within the entire range t = 0.14 to 0.39, the AUC achieved is significantly different from 0.5 with p-values all less than 0.008 (Mann-Whitney test) when T = 0.5.

For further discussion, we select t = 0.29 and T= 0.5 as this is within the longest continuous range of values with strong performance. However, it is important to point out that the performance of the integrated model is essentially the same for any value of t in this range (0.14 to 0.39). Figure 2 presents the AUC performance of IntMod for all values of the parameter t and for T = 0.5.

**Figure 2:**
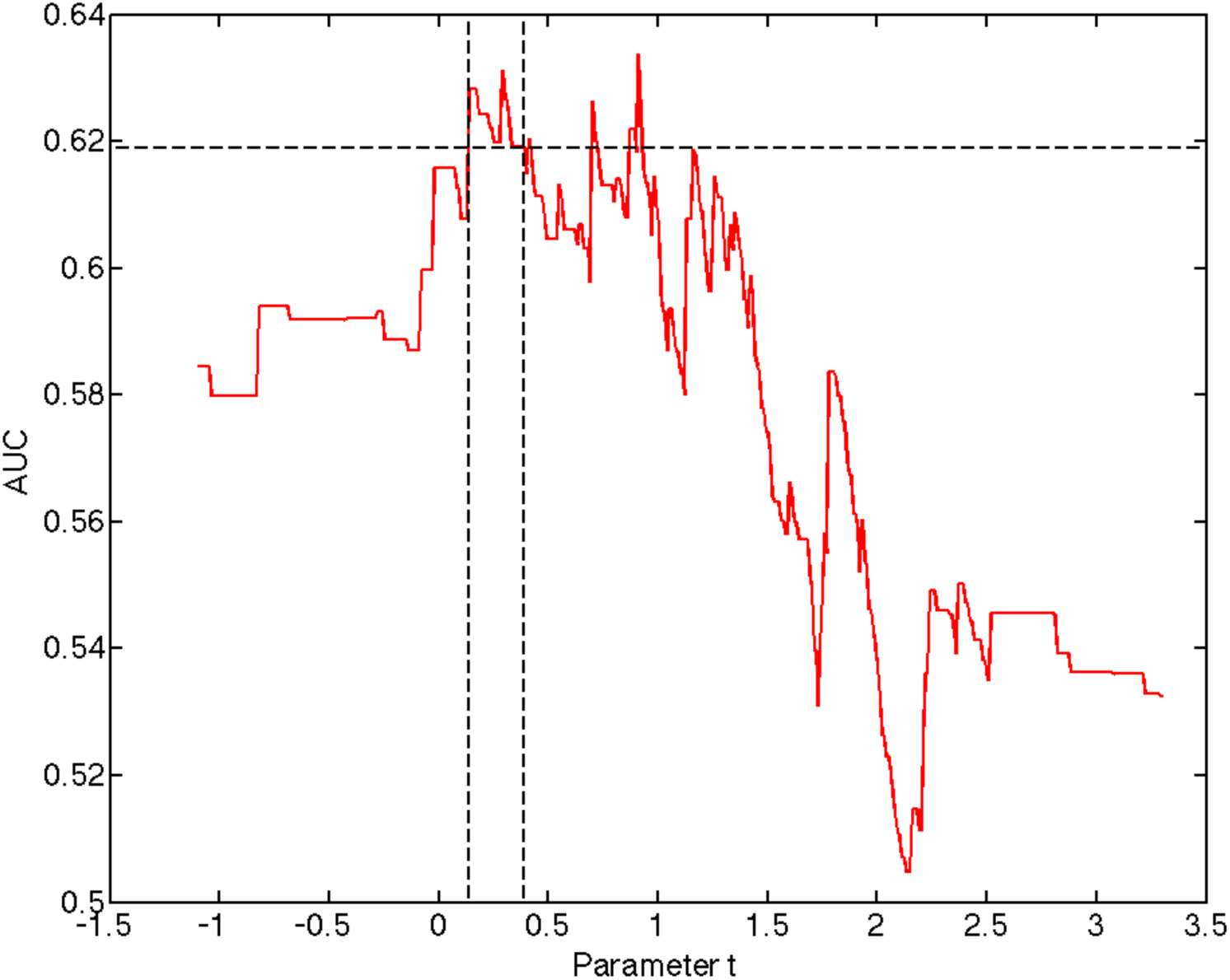
Performance of the Integrated Model (IntMod) for different values of parameter t and T= 0.5. Optimal sustained performance occurs for values of t between .14 and .39 where AUC values are all at least 62% and with p-values all below 0.008 (Mann-Whitney).

### Comparative Performance of the Proteomic Ratio, Mayo and the Integrated Model

On these subjects the AUC performance of the proteomic ratio, the simplified Mayo algorithm and the integrated model IntMod are, respectively, 60%, 58% and 63%. These are illustrated in Figure 3. Although these AUC values demonstrate an improved performance for the integrated model over the proteomic ratio and Mayo alone, we focus on the clinically relevant point on the IntMod ROC curve (sensitivity = 90%, specificity = 33%) to statistically compare performance at the same sensitivity or the same specificity. High sensitivity is typically required to rule out lung cancer confidently.

**Figure 3:**
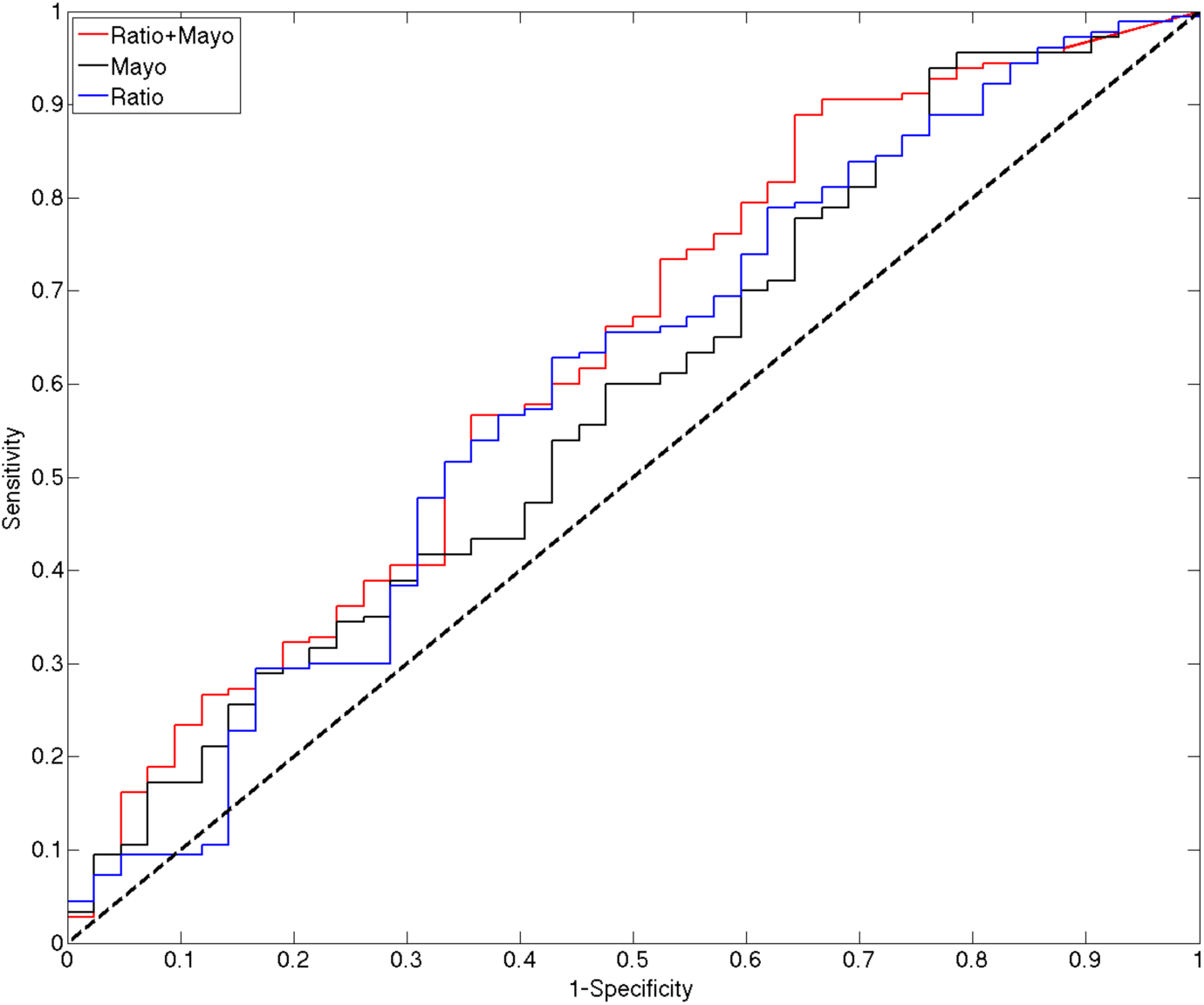
Comparison of proteomic ratio, the simplified Mayo algorithm and the Integrated Model (Ratio + Mayo. At sensitivity 90% and specificity 33% the integrated model has statistically significant better performance than both the simplified Mayo model and the proteomic ratio (see text for details).

Using the McNemar test, the integrated model has significantly better specificity (when sensitivity is fixed at 90%) to both the proteomic ratio (p-value 0.031) and the Mayo model (p-value 0.008). Similarly, using the McNemar test, the integrated model has significantly better sensitivity (when specificity is fixed at 33%) to both the proteomic ratio (p-value < 0.001) and the Mayo model (p-value < 0.001).

## Discussion

The two hypotheses tested in this study were confirmed. First, the proteomic marker LG3BP/C163A was more accurate than five commonly used clinical risk factors, alone or in combination, in the high prevalence population studied. Second, the integration of this proteomic marker with the five clinical risk factors resulted in a risk predictor statistically superior to both the proteomic marker and to the clinical risk factors separately. This result affirms that integrated risk predictors can enable better risk prediction for lung nodule management.

This analysis assessed five previously reported clinical risk factors and 11 proteomic markers on a new set of prospectively collected samples from 12 sites. The evaluation of risk markers on multiple sample sets helps to establish their performance and appropriate use in clinical use. In this study, we focused on the subpopulation of lung nodules between 8 and 20mm in diameter and having undergone an invasive procedure. This subpopulation is important as it contains benign lung nodules that are over-treated, and so, enables the evaluation of clinical risk factors and proteomic markers for the purpose of identifying those benign lung nodules that can avoid unnecessary invasive procedures.

The design of the integrated model warrants a few comments. First, proteomic ratios were assessed instead of individual protein markers. The reason for this is that ratios are particular useful as they normalize for pre-analytical and analytical variation (*14*), Furthermore, to reduce the possibility of overfitting we avoided utilizing more than two proteomic markers. Second, there are many other valid methodologies to integrate the clinical and proteomic risk factors including logistic regression (*17*), support vector machines (*18*) and random forests (*19*), among others, in addition to the decision tree approach used here.

To be clear, this is a discovery level study, using previously validated predictors to assess the above hypotheses. The integrated model developed needs to be validated in the intended use population, then assessed for clinical utility, prior to considering the clinical application of this biomarker. It is important to verify that tests shown to be accurate during clinical validation influence clinical decisions to the benefit of the patient. The biomarker reported here was designed as a rule out test (high sensitivity). The consequences of a true negative test in the population studied could be the avoidance of invasive testing in patients with benign nodules, while the consequences of a false negative test may be a delay in treatment of a malignant nodule. The proper clinical balance of these potential results is not clear, being influenced by available testing, patient characteristics and values. The actual clinical decisions that follow the test result will also be influenced by physician confidence in the result and understanding of its’ clinical application. As designed, a positive test result would not influence the result of the clinical risk prediction model used. The relatively low specificity of the test, due to the goal of ruling out malignancy, mandated this approach.

The integrated model was developed on a population of patients with lung nodules with a very high prevalence of lung cancer (>80%). This is in contrast to the average prevalence of lung cancer in nodules 8-20 mm in diameter (20-25%). The high prevalence suggests that physician judgement about proceeding with invasive testing in our cohort was quite good. We felt that it was important to develop this marker in a population where the etiology of the nodules was well established, leading to the choice of including only nodules that were biopsy confirmed. Ultimately, a rule out biomarker is likely to have greatest clinical utility in a population with a lower prevalence of lung cancer, perhaps in the 10-60% range. This intended use population will need to be targeted during clinical validation.

The clinical risk predictors, alone and in combination, did not perform as well as previously reported. This is related to the population studied. All patients had an invasive procedure performed. The clinical decision to perform a procedure reflects the physician’s judgement about the probability of malignancy in the nodule. As described in the results, there were no differences in the clinical and imaging variables included in the model between the lung cancer and benign nodule groups. Thus, the accuracy of the clinical risk models in the high prevalence population studied was poor. It will be important to confirm the hypotheses during clinical validation in the intended use population. Similarly, it will be important that any difference in accuracy leads to a clinically significant improvement in decisions and patient outcomes.

### Ethical approval and Consent

All procedures performed in studies involving human participants were in accordance with the ethical standards of the institutional and/or national research committee and with the 1964 Helsinki declaration and its later amendments or comparable ethical standards. Informed consent was obtained from all individual participants included in the study.

## Acknowledgements

Authors wish to acknowledge the efforts of participating sites and analysts at Integrated Diagnostics.

